# A tomato line hyposensitive to simulated proximity shade shows altered auxin-related gene expression and improved fruit yield under high-density field conditions

**DOI:** 10.1101/2025.03.18.643865

**Authors:** Esteban Burbano-Erazo, Silvana Francesca, Julia Palau-Rodriguez, Andrea Berdonces, Laura Valverde-Carbajal, Jose Perez-Beser, Matteo Addonizio, Jaume F. Martinez-Garcia, Maria Manuela Rigano, Manuel Rodriguez-Concepcion

**Affiliations:** Institute for Plant Molecular and Cell Biology (IBMCP), CSIC-Universitat Politècnica de València, 46022 València, Spain; Department of Agricultural Sciences, University of Naples Federico II, 80055 Portici (Naples), Italy; Doctorate in Biotechnology, Facultat de Farmàcia i Ciències de l’Alimentació, Universitat de Barcelona, Avda. Diagonal 643, 08028 Barcelona, Spain

**Keywords:** auxin, field, fruit, shade, tomato, yield

## Abstract

Plants detect the presence of nearby vegetation as a reduction in the ratio of red to far-red light (low R/FR). This proximity shade signal can be simulated in the lab by supplementing white light (W) with FR (W+FR). While shade avoidance strategies are considered undesirable in agricultural crops, FR supplementation enhances plant growth and fruit quality in tomato (*Solanum lycopersicum*).

Here we compared the response of different tomato genotypes to W+FR in the lab and identified one *S. pennellii* introgression line (IL2-2) with a shade-tolerant phenotype at the seedling stage.

Compared to the shade-avoider parental genotype M82, IL2-2 plants showed reduced elongation upon W+FR exposure and a disrupted expression of auxin-related genes both under W and W+FR. At harvest, W+FR treatment improved M82 fruit quality by increasing °Brix, ascorbic acid and carotenoids, and these quality traits remained virtually unchanged in IL2-2. Under high density (HD) conditions, fruit quality traits were hardly impacted by planting density or genotype, but IL2-2 showed improved fruit yield.

Our findings suggest that IL2-2 could serve as a valuable genotype for high-density or intercropping agrosystems.

## Introduction

Plants use light not only as an essential source of energy for photosynthesis but also as a signal informing on the presence of nearby vegetation that can potentially become competitors. Leaves and other photosynthetic (i.e. green) tissues absorb red light (R) but not far-red light (FR). As a consequence, light reflected from the leaves of neighboring plants has a much lower R/FR ratio than direct sunlight. This decrease in R/FR is a signal of vegetation proximity that does not require actual shading or reduced light intensity. The low R/FR signal (herein referred to as proximity shade) is perceived by the phytochrome family of photoreceptors, particularly phytochrome B (phyB). After phyB-dependent perception, the signal is transduced by a network of PHYTOCHROME INTERACTING FACTORs (PIFs), a group of transcription factors that function as inducers or repressors of the proximity shade response (Casal & Fankhauser, 2023; Martinez-García & Rodriguez-Concepcion, 2023). In the lab, proximity shade can be simulated by enriching white light (W) with FR, hence reducing the R/FR ratio without changing total light intensity. This simulated shade (W+FR) treatment triggers elongation growth and degradation of photosynthetic pigments (chlorophylls and carotenoids) in plants that are shade-intolerant, including *Arabidopsis thaliana*, tomato (*Solanum lycopersicum*), and most crops (Cagnola et al., 2012; Bou-Torrent et al., 2015; Chitwood et al., 2015; Llorente et al., 2016; Ortiz-Alcaide et al., 2019; Fanwoua et al., 2019; Ji et al., 2019; Kim et al., 2019; Ji et al., 2020; Sun et al., 2020; Ji et al., 2021; Casal & Fankhauser, 2023; Li et al., 2024; Shomali et al., 2024). The response to proximity shade, referred to as shade avoidance syndrome (SAS), is strongly attenuated in shade-tolerant species such as *Cardamine hirsuta*, an edible relative of *Arabidopsis*. Exposure of *Cardamine* plants to W+FR causes no elongation growth and it only leads to a minor decrease in the levels of chlorophylls and carotenoids compared to W-grown controls (Molina-Contreras et al., 2019; Morelli et al., 2021).

Domestication has led to increased tolerance to proximity shade as the SAS involves responses that are not desirable from an agronomic perspective, such as excessive elongation growth at the expense of defense against biotic stress. However, the idea that attenuating shade-avoidance responses would benefit crops might be an oversimplification (Gommers et al., 2023; Casal & Fankhauser, 2023). Agricultural practices directly impacted by proximity shade, such as high density planting or intercropping, also involve additional light-independent competence for access to water and nutrients and increased risk of infections associated with crowded environments. Besides, it remains to be demonstrated that the SAS mechanisms deduced from studies using FR supplementation to simulate shade also apply to actual plant proximity shade in the field. In tomato, a number of recent studies have observed positive effects of FR supplementation in plant growth but also in fruit quantity (number and weight) and quality (sugar content) (Fanwoua et al., 2019; Ji et al., 2019; Kalaitzoglou et al., 2019; Ji et al., 2020). However, it is unclear whether these conclusions could also apply to agricultural settings involving proximity shade with no FR supplementation. For example, high density conditions are associated with low R/FR but they also involve reduced light intensity in the lower strata of the crop plants and increased competence for resources.

Here we investigated the variability in the response to simulated shade in tomato during short exposure to W+FR at early stages of development (e.g., in seedlings) and long-term exposure up to the fruit-bearing stage. We further identified a *S. pennellii* introgression line showing hyposensitivity to proximity shade in terms of growth and a gene expression profile consistent with a heavily impaired homeostasis of auxin, the main hormone controlling proximity shade-induced elongation. Furthermore, we compared how FR supplementation in the greenhouse and high density conditions in the field influenced long-term plant growth and fruit traits of this shade-tolerant line and the shade-avoider parental line.

## Materials and Methods

### Plant material and treatments

A protocol was set up on plants at seedling stage to investigate the response to proximity shade in tomato (*Solanum lycopersicum* L.) genotypes Money Maker, San Marzano, M82 and MicroTom, performing at least 4 independent experiments. Additionally, 2 populations of tomato landraces and introgression lines (IL) were evaluated. In particular, we used 21 landraces (8 from South America, 4 from USA, 9 from other European and world regions), and on 20 *Solanum pennellii* introgression lines (IL) belonging to a collection available at the University of Naples Federico II, Department of Agricultural Sciences (details hosted in the LabArchive repository http://dx.doi.org/10.6070/H4TT4NXN). For all the genotypes, seeds were sterilized and plated on solid MS 0.5x MS medium without added vitamins containing 1% (w/v) agar (Barja et al., 2021). Plates were incubated in the dark for 4 days. Only seeds germinated at the same time were transferred to soil and kept for 2 days in a walk-in growth chamber under a photoperiod of 8 h of darkness and 16 h of white light (W) at a photosynthetic photon flux density (PPFD) of 150 ± 20 μmol m^-2^ s^-1^ at 25 ± 1 °C. W was obtained using a mix of 5 Phillips MAS LEDtube 1500mm HO 23W840 and 4 MAS LEDtube 1500mm HO 23W830 LED tubes, arranged in an alternating pattern. To simulate proximity shade, GreenPower LED module HF far-red (Philips) tubes supplying FR were added between the racks of LED tubes supplying W. As a result, in the same chamber there was an area illuminated with W (R/FR of 3.14) and another area illuminated with W+FR (simulated shade, R/FR of 0.17). W and W+FR areas with the same PPFD (ca. 150 μmol m^-2^ s^-1^) were separated by an opaque panel so W-grown plants were not exposed to residual FR. After the 2-day adaptation period under W, seedlings were either left under W or transferred to W+FR. Eleven days after the treatment, morphological measurements were done using the IMAGE J software (Schneider et al., 2012). MicroTom plants were also grown in the same growth chamber for 4 months, until the fruit-bearing stage. In the case of IL2-2 plants and the M82 parental, growth beyond the seedling stage was carried out in a greenhouse room equipped with W and W+FR light sources to support the same photoperiod and light intensity (PPDF) and quality (R/FR ratio) values similar to those in the growth chamber. Seeds were sown in seed trays and after germination the seedlings were transferred to plastic pots (21 cm diameter) with commercial substrate. Plants were grown in the greenhouse room with 24 ± 3 ◦C air temperature during the day and 18 ± 3 ◦C during the night. Plants were irrigated using the following full nutrient solution: Ca(NO₃)₂ 1.27 g L^−1^ ; P2O5 0.26 g L^−1^ ; K2O 0.23 g L^−1^; MgSO₄ 0.31 g L^−1^. The electrical conductivity (EC) of the NS was 1.8 ± 0.1 dS m^−1^. The pH was monitored daily and maintained at 6.0 ± 0.3. All plants were grown under W (PPDF of 190 ± 15 μmol m^-2^ s^-1^) for two weeks. Thereafter, the plants were subjected to two different light treatments using a custom made lighting system developed by MEG s.r.l (Milan, Italy). As previously outlined, the light treatment comprised enrichment with FR light, maintaining the same flux density (as the W control). Consequently, only the R:FR was reduced to 0.3 (W+FR). The analysis of morphological, physiological and molecular parameters was conducted on plants that had reached the thirty-day growth stage and at the end of the reproductive cycle.

*Arabidopsis thaliana* Col-0, *sav3-5* and *pif457* mutants have been described before (Pastor-Andreu et al., 2024). For hypocotyl assays, seeds were surface-sterilized and sown on solid growth medium without sucrose (0.5xMS-) at two different densities: low (2 seeds·cm^-2^) or high (18 seeds·cm^-2^). After stratification (3-4 days in the dark at 4°C), plates with seeds were incubated in dedicated growth chambers (Aralab, model D1200PL) at 22°C and continuous W emitted from LED light tubes that provided a PPFD of 55 μmol m^-2^ s^-1^ (R/FR of 3.55). Hypocotyl length was measured as indicated elsewhere (Urdín-Bravo et al., 2024). Three biological replicates were measured; each consisting of at least 15 seedlings per treatment (density).

### Open field experiment

The field trials were carried out in experimental farms (ARCA2010 in Acerra, Italy, and Fundación Cajamar in Paiporta, Spain). At the end of April 2024, three weeks after sowing, when the third true leaf was fully expanded, tomato plants (genotypes M82 and IL2-2) were transplanted in the open field. The experiment was designed to assess the response of tomato yield components to high-crop density (HD) compared to normal farmer practice (low-crop density; LD). The distance from plant to plant in LD and HD was 33 cm and 16 cm, respectively (Supplemental Figure S3). Plants were arranged in a randomized block design with four replications per treatment and eight plants per biological replication. The experimental field was irrigated with a drip irrigation system of 4 L h^-1^ (one emitter per plant). The irrigation and nutrition of the plants was done according to the management practices of the area.

### Assessment of pigment content

The leaf pigment index and chlorophyll content parameters were determined on five fully expanded leaves per genotype per treatment using a portable meter SPAD-502 Plus (Konica Minolta Optics, Japan). Leaf chlorophylls and carotenoids were also quantified by HPLC-DAD (Barja et al., 2021).

### Photosynthetic parameters

Gas exchange and chlorophyll fluorescence measurements were conducted on five healthy, young, fully expanded leaves (typically the third leaf counting from the apical meristem) of adult plants of each genotype per treatment. Leaf gas exchanges were measured at 30 days after transplant with a portable photosynthesis system (LCA 4; ADC BioScientific Ltd., Hoddesdon, UK) equipped with a broadleaf chamber (cuvette window area, 6.25 cm^2^). All the measurements were conducted in an open system at a temperature of 24±0.5°C, a CO2 concentration of 400±10 μmol m^−2^ s^−1^, and a relative humidity of 70 – 80%. The measurements were carried out with a transparent top leaf chamber, allowing measurements at ambient light intensity while including or excluding FR light. From gas-exchange measurement the following instantaneous parameters were measured: sub-stomatal CO2 (vpm), transpiration (E; mmol m^−2^ s^−1^), stomatal conductance (gs; mol m^−2^ s^−1^), photosynthetic rate (μmol m^−2^ s^−1^).

### Fruit traits

Fruits were quantified and weighted after harvest, before collecting pericarp samples (that were snap-frozen in liquid nitrogen for storage) and counting seed number. Fruit equatorial area was calculated using the IMAGE J software from pictures taken from above to harvested fruits. Total solid soluble content (°Brix) was measured with a refractometer (Hanna Instruments, Padova, Italy). Fruit hardness, expressed as the maximum force (kg cm^-2^) needed for penetration of the probe (8 mm diameter) into the fruit through the tomato skin, was measured using a penetrometer (PCE-PTR200 penetrometer, Capannori, Italy). Pericarp tissue was used to measure ascorbic acid by using a colorimetric method (Francesca et al., 2022) and carotenoids by HPLC-DAD (Barja et al., 2021).

### RNA sequencing

Total RNA was extracted from the first two completely expanded leaves of 2-week-old plants exposed to W+FR (or W as a control) for 24 h. The tissue was collected and immediately frozen in liquid nitrogen for storage at -80 °C. RNA extraction was carried out using the RNeasy Mini Kit (Qiagen, Hilden, Germany) according to the method reported by the manufacturer. Three independent replicates of total RNA (1 µg) extracted from leaves of different plants growing under the same conditions was used for RNA sequencing by BGI Tech Solutions (https://services.bgi.com). A total of 12 samples were tested using the DNBSEQ platform, with an average yield of 6.68G data per sample. The average alignment ratio of the sample comparison genome (*S. lycopersicum* 4081.JGI.ITAG2.4.v2201) was 97.82%. The default DrTom pipeline was used to estimate gene expression, identify DEGs, and analyze enrichment according to GO terms or KEGG pathways.

### RT-qPCR analyses

Total RNA from frozen leaf samples was isolated using the Purelink RNA mini kit (Thermo Fisher Scientific, Waltham MA, USA). Determination of RNA integrity and quantity as well as cDNA synthesis were carried out as described (Ezquerro et al., 2023). Transcript abundance of *YUC9* (Solyc06g083700) and *PIF7b* (Solyc06g069600) was assessed using the following gene-specific primers (5’ – 3’): YUC9_qF, ACCGTTGAACTTGTCACTGG; YUC9_qR, GCACCAGCTAGCCCTTTCC; PIF7b_qF, GCGCTCCCCTCATTTATCCA; and PIF7b_qR, TTTGGGGCTGATGGATTCGG. The tomato gene Solyc07g025390 was used as endogenous reference gene for normalization (Gonzalez-Aguilera et al., 2016). The RT-qPCR was carried out on a QuantStudio 3 Real-Time PCR System (Thermo Fisher Scientific) using three independent samples and two technical replicates of each sample.

### Statistical analyses

Statistical analyses, including t-test, one-way and two-way ANOVA with means comparison tests were carried out using the statistical software R v4.1.0 (https://www.R-project.org/).

## Results

### Tomato genotypes show a range of responses to simulated proximity shade (W+FR)

Tomato plants of different accessions (Money Maker, San Marzano, M82 and MicroTom) were germinated in darkness and evenly grown 4-day-old seedlings were transferred to soil and incubated under W for 2 additional days. Then, half of them were transferred to W+FR (referred to as simulated shade) and the other half were left under W for 11 days. At the end of the treatment, different parameters associated to the SAS were measured (Figure 1A).

**Figure 1.**
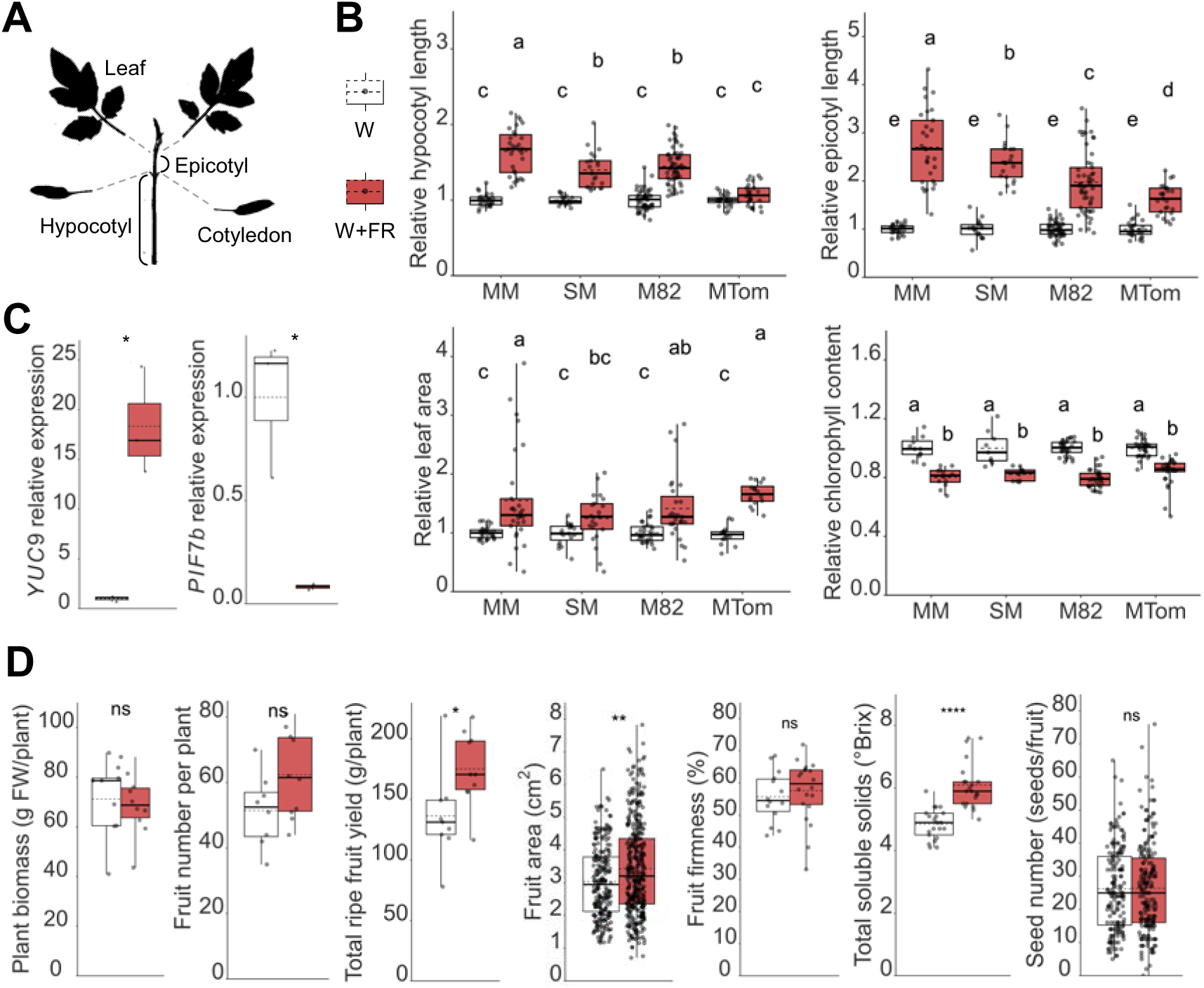
Tomato accessions show a range of responses to simulated shade. (A) Schematic representation of tomato seedling organs used to quantify the impact of simulated shade. (B) Effect of simulated shade in different organs of tomato seedlings of the MoneyMaker (MM), San Marzano (SM), M82 and MicroTom (MTom) accessions germinated under darkness for 4 days and grown under white light (W) for 2 days and then exposed to FR-supplemented W (W+FR) or left under W for 11 days. (C) RT-qPCR analysis of shade marker genes *YUC9* and *PIF7b* in leaves from W-grown 2-week-old MicroTom seedlings exposed to simulated shade (W+FR) or left under W for 24 h. (D) Quantification of total biomass and fruit-associated features of MicroTom plants grown under W or W+FR for 4 months. In all the plots, dots represent individual data points. The lower boundary of the boxes indicates the 25th percentile, the black line within the boxes marks the median, the dotted line within the boxes marks the mean, and the upper boundary of the boxes indicates the 75th percentile of the data distribution. Whiskers above and below the boxes indicate the minimum and maximum values (excluding outliers). Statistically significant differences are represented with letters (one-way ANOVA followed by Duncan’s multiple range test, P < 0.05) or asterisks (t-test, ns: not significant; *: P < 0.05; **: P < 0.01; ****: P < 0.001).

All accessions tested showed longer epicotyls and lower leaf chlorophyll contents when exposed to W+FR (Figure 1B). Leaf area also increased under simulated shade, even though the difference with W controls was not statistically significant in San Marzano (Figure 1B). In the case of hypocotyl elongation, by far the most studied SAS phenotype in *Arabidopsis* (Martinez-García & Rodriguez-Concepcion, 2023), only MicroTom plants failed to respond to the W+FR treatment (Figure 1B). To check whether MicroTom plants responded to the W+FR treatment at the molecular level, we used marker genes shown to be up-regulated (*YUC9*) or down-regulated (*PIF7b*) by low R/FR in seedlings of the tomato Ailsa Craig cultivar (Sun et al., 2020). RT-qPCR analysis showed a strong *YUC9* up-regulation and *PIF7b* down-regulation in leaves of MicroTom seedlings exposed for 24h to W+FR compared to W controls (Figure 1C), confirming that MicroTom plants perceived and transduced the low R/FR signal.

To next investigate long term responses to proximity shade, the MicroTom accession was selected based on its small size, which allowed to grow plants and estimate fruit yield in the growth chamber. In this case, plants were germinated and grown for 4 days as described above and then they were left under W or transferred to W+FR for 4 months. FR supplementation increased fruit yield, likely as a combined result of very slight increases in fruit size and number, with no differences detected in plant weight (Figure 1D). Ripe fruit from plants grown under W+FR showed an improved content of total soluble solid (°Brix) compared to W controls, but no differences were detected in hardness or in seed number per fruit (Figure 1D). The enhancement of fruit production and °Brix has been previously associated to FR-supplemented conditions in several tomato cultivars (Fanwoua et al., 2019; Ji et al., 2019; Kalaitzoglou et al., 2019; Ji et al., 2020), confirming that both short-term and long-term responses to low R/FR are conserved in different tomato accessions, including MicroTom.

### Screening for shade-tolerant tomato lines

We next used the experimental conditions optimized for seedlings to screen for tomato lines showing attenuated responses to proximity shade. To increase genetic variability, we used collections of introgression lines (ILs) and landraces (Figure 2 and Supplemental Figure S1). In particular, twenty ILs covering the genome of the wild tomato relative *S. pennellii* in the genetic background of the tomato (*S. lycopersicum*) cultivar M82 (Eshed & Zamir, 1995) and twenty-one landraces from Europe, South America, USA, and other world regions (Ruggieri et al., 2014), were included in the screening for shade-hyposensitive lines. As a read-out, we used epicotyl elongation under W+FR compared to W, as it was the phenotypic trait showing the strongest response and dynamic range (Figure 1).

**Figure 2.**
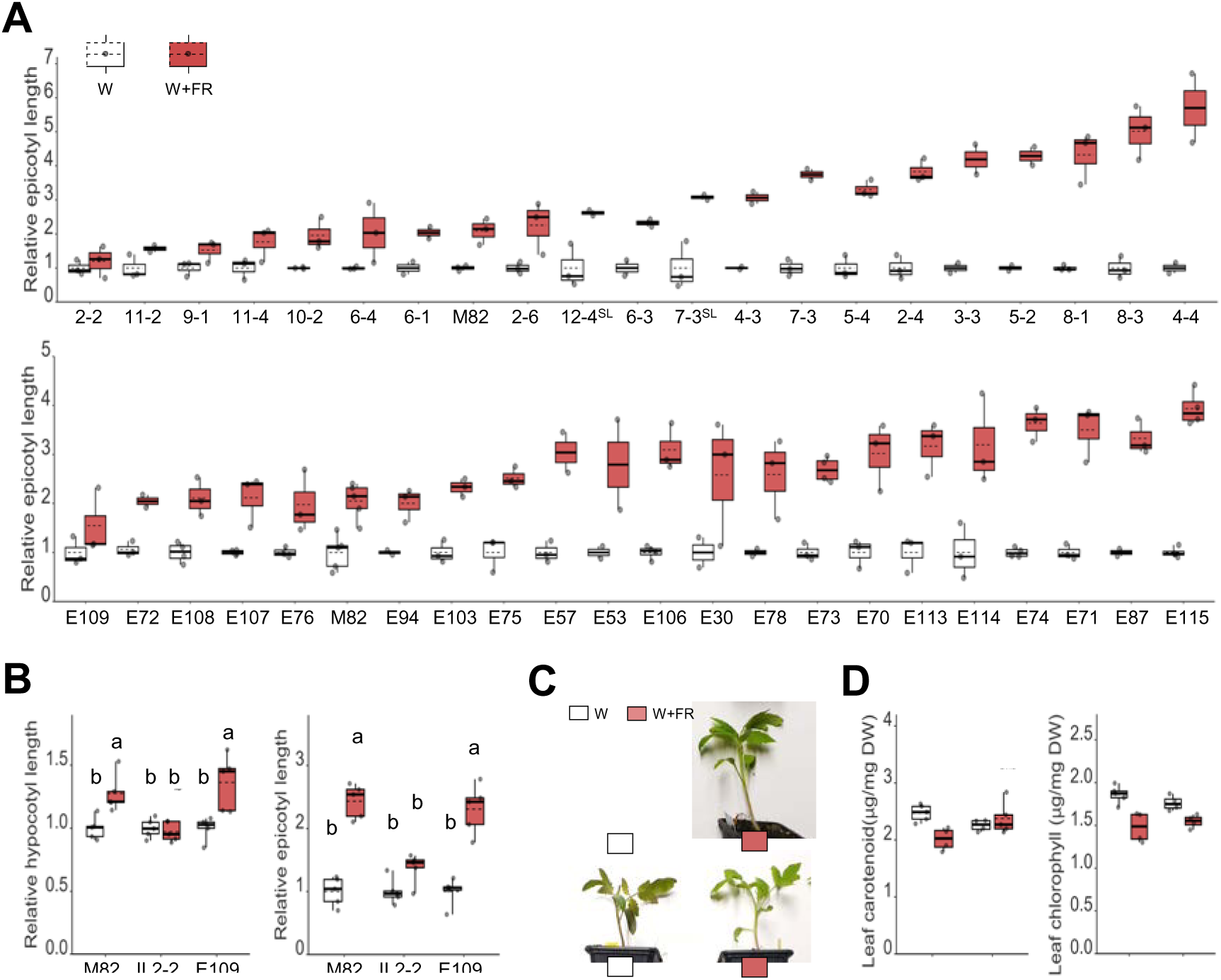
The tomato introgression line IL2-2 shows an attenuated response to simulated shade. Seeds from different tomato lines were germinated and grown under white light (W) and then exposed to FR-supplemented W (W+FR) or left under W for 11 days. (A) Epicotyl elongation of the indicated tomato introgression lines (ILs, upper plot) and landraces (lower plot) in response to simulated shade. SL, sub-line. Values are shown relative to W controls. (B) Hypocotyl and epicotyl length of the indicated lines represented as described in (B). (C) Representative images of seedlings from the IL2-2 line and its M82 parental. (D) Levels of photosynthetic pigments in the indicated seedlings. Boxplot legend and statistical analysis were as described in Figure 1 legend.

Most ILs and landraces were found to display an exacerbated epicotyl elongation response to low R/FR compared to M82 (Figure 2A). Among the few lines that showed a reduced elongation response to W+FR, IL2-2 and E109 seedlings showed the strongest phenotype and a similar epicotyl length under W compared to M82 (Figure 2A and Supplemental Figure S1). To validate the shade-hyposensitive phenotype of these two lines, a second round of experiments was performed using only lines M82, IL2-2 and E109. The results confirmed that IL2-2 seedlings showed a significantly reduced response to simulated shade compared to the M82 parental line not only in terms of epicotyl elongation but also when measuring the increase in hypocotyl length (Figure 2B). In the case of the E109 line, however, the elongation response of hypocotyls and epicotyls was similar to that of M82 seedlings (Figure 2B). We therefore concluded that only the IL2-2 line showed an attenuated elongation response to simulated shade (Figure 2C).

To test whether other short-term responses to low R/FR were also attenuated in IL2-2, we quantified the levels of photosynthetic pigments (carotenoids and chlorophylls) by HPLC after simulated shade exposure (Figure 2D). M82 and IL2-2 seedlings germinated and grown in plates were transferred to soil and left under W before exposing them to W+FR (or left under W) for 24 h. Treatment with W+FR caused a reduction in the levels of carotenoids and chlorophylls in both M82 and IL2-2 seedlings, but the reduction appeared to be slightly attenuated in the case of the introgression line (Figure 2D).

### Auxin-related genes are misregulated in IL2-2 plants and respond less to W+FR treatment

The response of M82 and IL2-2 lines to simulated shade was next compared at the transcriptomic level. Dark-germinated M82 and IL2-2 seedlings were transferred to soil at day four after imbibition and grown under W for 11 days. Then, some of them were left under W and others were exposed to W+FR for 24h before extracting RNA from leaves. Following RNA sequencing (RNA-seq) and bioinformatic analyses, genes with more than 2-fold change (FC) in expression (log2FC>1, P-value < 0.05) were considered differentially expressed genes (DEGs) (Figure 3). The reduced morphological and metabolic responses to simulated shade displayed by IL2-2 seedlings (Figure 2) appeared in contrast with a substantial increase in number of genes differentially expressed in response to the W+FR treatment compared to M82 (Figure 3A). GO Biological Process (GO-P) enrichment analyses showed that the 292 genes responding to simulated shade in M82 but not in IL2-2 seedlings were involved in light-regulated processes related to growth, including auxin signaling, photomorphogenesis, phototropism, and cell wall biogenesis (Figure 3B). Consistent with the central relevance of auxin for shade-promoted growth in tomato (Cagnola et al., 2012; Bush et al., 2015; Sun et al., 2020), the highest enrichment Q-value was found for “auxin-activated signaling pathway” genes, which encode proteins required for auxin transport (PIN3, PIN4, BG1LA, BG1LE) and response (IAA1, IAA7, IAA17, IAA22, ARF10B) (Figure 3C). Furthermore, four auxin signaling genes encoding SAUR and ATHB2 homologs were found among the top-20 genes most up-regulated by simulated shade in M82 plants (Figure 3D). Most of these auxin-related genes were upregulated in IL2-2 compared to M82 in plants grown under W but they showed similar levels in M82 and IL2-2 plants after the W+FR treatment (Figure 3C-D).

**Figure 3.**
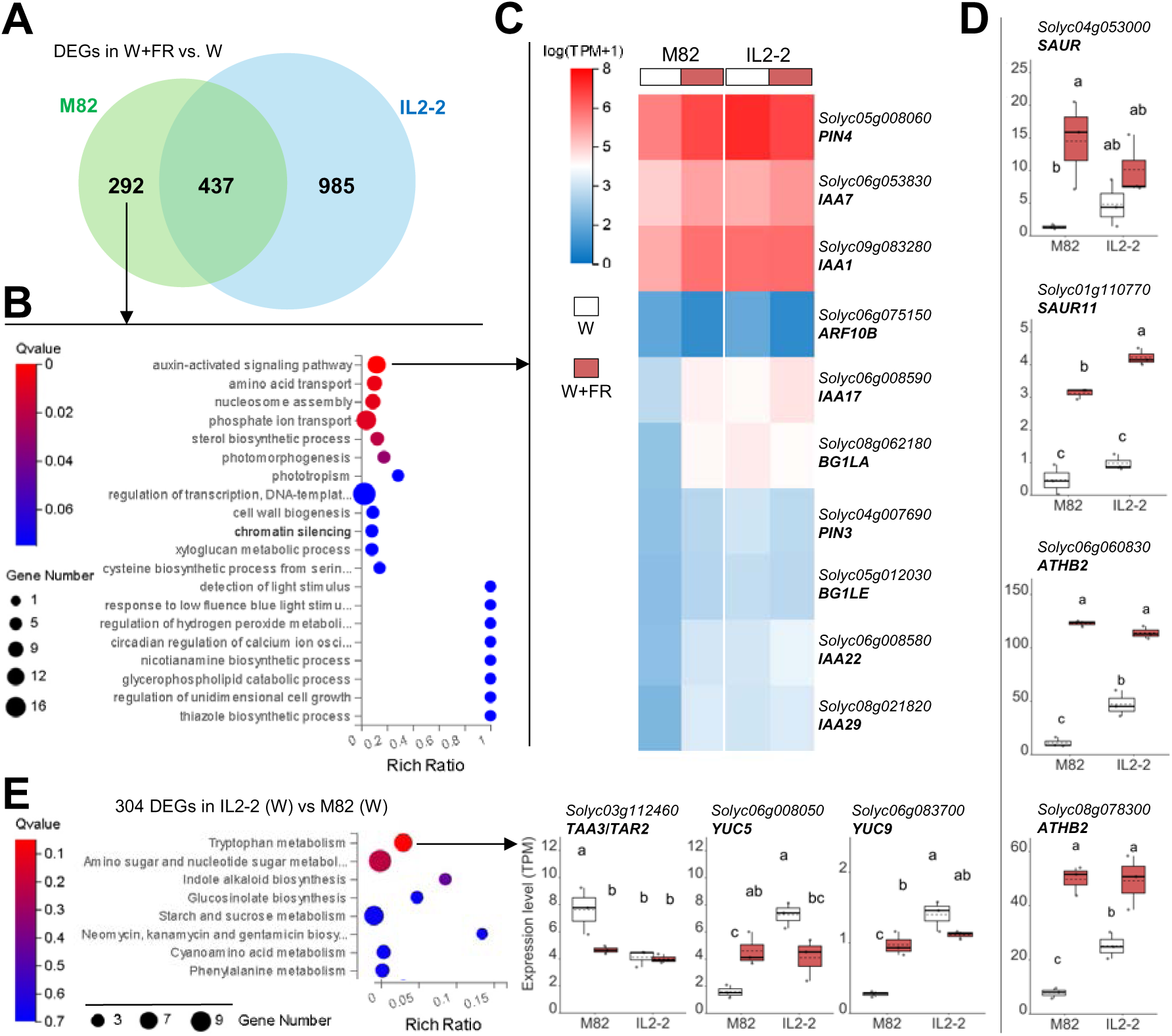
IL2-2 plants show an altered expression of auxin-related genes. Two-week-old M82 and IL2-2 seedlings were treated with W+FR (or W as a control) for 24 h and then used for RNA sequencing. (A) Venn diagram of differentially expressed genes (DEGs) in plants exposed to simulated shade. (B) GO term (Biological Process) enrichment analyses of the 292 DEGs specific of M82. The phyper function in R software was used to calculate the enrichment (Rich) ratio and the Q-value, which represents the significance value of enrichment (Q-values lower than 0.05 correspond to significant enrichment). The size of the bubbles represents the number of DEGs annotated to a particular GO term. (C) Heatmap comparison of transcript abundance (mean log2 TPM+1) for the genes included in the GO term “auxin-activated signaling pathway”. (D) Expression of the 4 auxin signaling genes found among the top 20 most highly up-regulated DEGs in simulated shade-treated M82 plants. TPM (Transcripts Per Kilobase Million) values are represented. (E) KEGG pathway enrichment analyses of the DEGs found between M82 and IL2-2 seedlings grown under control (W) conditions and transcript levels of the genes included in the term “tryptophan metabolism”. Bubble plot and box plots were made as described in (B) and D). Boxplot legend and statistical analysis were as described in Figure 1 legend.

GO-P analysis of the 304 genes that were differentially expressed in M82 and IL2-2 seedlings grown under control (W) conditions showed the second highest Q-value for the same category of auxin-activated signaling pathway genes that was overrepresented in shade-regulated M82 genes. A different enrichment analysis of the same 304 DEGs using KEGG pathways showed the highest Q-value for tryptophan metabolism, a category that included genes for auxin biosynthetic enzymes such as TAA3/TAR2, YUC5 and YUC9 (Figure 3E). Again, these genes were misregulated in IL2-2 and reached similar expression levels as the M82 parental following exposure to simulated shade. Also similar to the auxin signaling and transport genes, biosynthetic genes were less responsive or responded differently to W+FR in IL2-2 seedlings, likely contributing to the reduced shade-triggered elongation responses observed in the introgression line (Figure 2). In the case of genes involved in photosynthetic pigment biosynthesis, M82 and IL2-2 seedlings showed a much more similar profile both under W and in response to the W+FR treatment (Supplemental Figure S2).

### The shade-hyposensitive phenotype of IL2-2 does not alter fruit quality under FR-supplemented light

To analyze long-term responses to simulated shade in IL2-2 plants, they were grown together with the parental line M82 in a greenhouse room that, similar to the growth chamber used to this point, had separate areas illuminated with W or W+FR. Following germination in seed trays, M82 and IL2-2 seedlings were transferred to plastic pots. After two weeks under W in the greenhouse room, half of the pots were left under W and the other half exposed to W+FR. Similar to that observed in the growth chamber with 15-day-old seedlings that had been exposed for 11 days to either W or W+FR (Figure 2C), 30-day-old M82 plants growing in the greenhouse room and exposed for 10 days to either W or W+FR were taller and paler under simulated shade (Figure 4A). This result confirmed that the low R/FR conditions in the greenhouse room were working to activate the response to proximity shade. In the case of IL2-2 plants, their height was virtually unaffected by the FR supplementation of W light (Figure 4B). While they were also slightly paler under W+FR, the shade-induced chlorophyll drop was not significantly attenuated in IL2-2 plants compared to M82 controls (Figure 4B). When plants produced fruit, we compared both the vegetative and the reproductive (fruit) phenotypes under W or W+FR. Continuous growth under W+FR resulted in longer plants in the case of M82, whereas no significant difference in plant height was observed for the IL2-2 line growing under W+FR compared to control W conditions (Figure 4C). While total plant biomass was unaffected by the treatment or the genotype, W+FR treatment resulted in a higher photosynthetic rate in M82, an increase that was attenuated in the IL2-2 line (Figure 4C). In both genotypes, FR supplementation elevated transpiration rate (E) and reduced the substomatal CO₂ concentration, without changing the stomatal conductance rate (gs) (Figure 4C). Fruit production was very low in our greenhouse conditions (less than a dozen fruits per plant), making it impossible to reliably calculate yield. As for fruit features, M82 plants grown under W+FR conditions produced fruit with higher soluble solid content (°Brix) but only marginal differences in area (size) or hardness compared to W controls (Figure 4D), mostly paralleling that observed for the MicroTom accession (Figure 1D). Also similarly, seed number per fruit was unaffected by FR supplementation (Figure 4D). A deeper analysis of M82 ripe fruit showed that W+FR treatment resulted in increased accumulation of reduced ascorbate (vitamin C) and carotenoids (pro-vitamin A), hence adding nutritional quality to the tomatoes (Figure 4D). A very similar fruit and seed phenotype was observed for the IL2-2 line, indicating that its shade hyposensitivity mostly impacts plant vegetative growth (height) but does not prevent the positive effects on fruit quality provided by FR supplementation.

**Figure 4.**
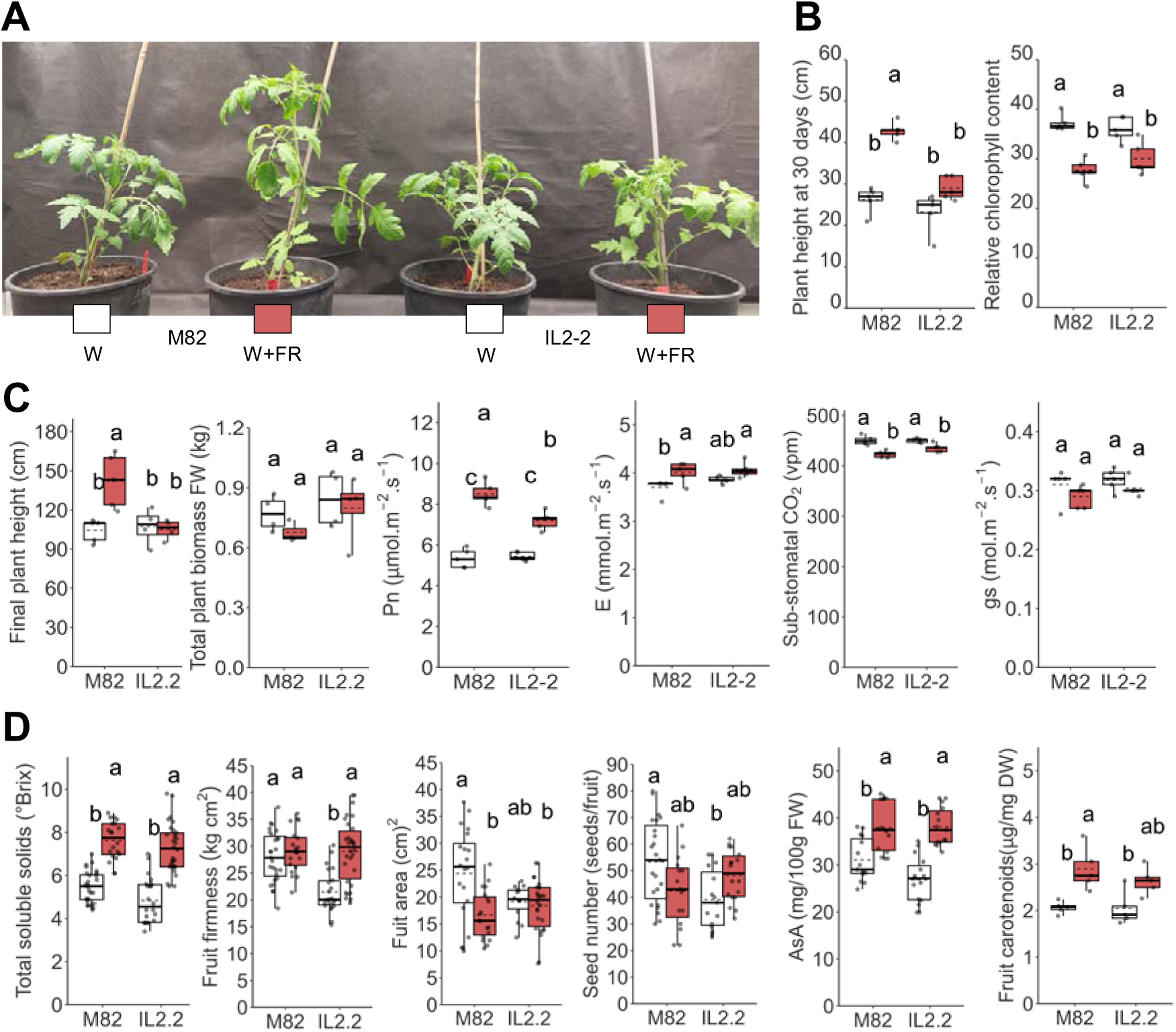
IL2-2 plants elongate less than M82 but show a similar enhancement of fruit quality under FR-supplemented light. (A) Representative images of M82 and IL2-2 plants germinated and grown under W for 20 days and then transferred to W+FR or left under W for 10 more days. (B) Plant height and chlorophyll content of plants like those shown in (A). (C) Plant height, biomass and photosynthetic parameters of M82 and IL2-2 plants kept under W or W+FR until fruit production. (D) Quantification of fruit-associated features from the plants described in (C). Boxplot legend and statistical analysis were as described in Figure 1 legend.

### IL2-2 plants show an improved fruit yield under high-density conditions in the field

We next aimed to test whether some of the conclusions derived from the results using simulated shade (W+FR) and pot-grown plants in the greenhouse could also apply to soil-grown plants in open field conditions exposed to a natural proximity shade provided by neighboring plants. As IL2-2 plants showed a strong alteration of auxin-related processes (Figure 3), we first examined the relevance of this hormone for enhanced growth under higher density conditions (Figure 5).

**Figure 5.**
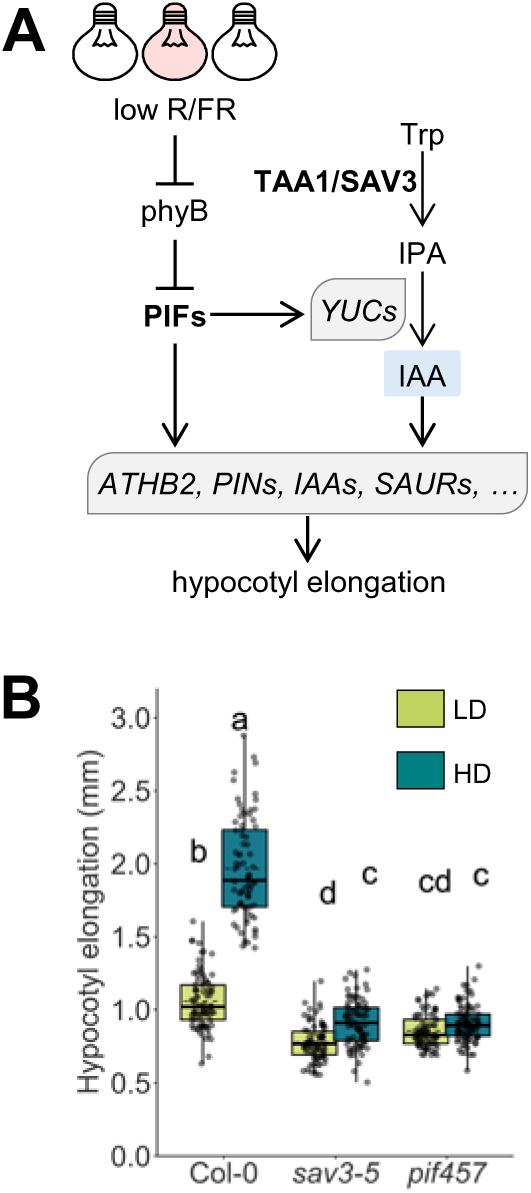
Auxin is required for hypocotyl elongation under high planting densities. (A) Cartoon depicting the main proximity shade signaling components and their connection with auxin production and hypocotyl growth. (B) Arabidopsis *sav3-5* and *pif457* mutant lines and their wild type parental (Col-0) were plated either at low-density (LD) or high-density (HD) and grown for one week under continuous W. Dots represent measurements of hypocotyl length in individual seedlings growing in three different plates. Boxplot legend and statistical analysis were as described in Figure 1 legend.

For this purpose, we employed *Arabidopsis* plants, in which a direct link between low R/FR perception by phyB and PIF-regulated auxin production has been well established (Casal & Fankhauser, 2023; Martinez-García & Rodriguez-Concepcion, 2023). In this pathway, tryptophan (Trp) is converted to indole-3-pyruvic acid (IPA) by TRYPTOPHAN AMINOTRANSFERASE OF ARABIDOPSIS 1 (TAA1), also known as SHADE AVOIDANCE 3 (SAV3), and then flavin monooxygenases of the YUCCA (YUC) subfamily catalyze the oxidative decarboxylation of IPA to indole-3-acetic acid (IAA), the major natural auxin (Figure 5A). PIF4, PIF5 and PIF7 are the three main PIFs inducing the expression of *YUC* genes, and they also regulate the expression of other genes involved in auxin homeostasis, transport, and signaling, eventually promoting hypocotyl elongation (Figure 5A). Mutants deficient in SAV3 (*sav3-5*) or in PIF4, PIF5 and PIF7 (*pif457*) are therefore defective in IAA synthesis and hypocotyl elongation in response to W+FR (Pastor-Andreu et al, 2024). To check whether this auxin-dependent pathway was also important to regulate growth in response to plant density conditions, *sav3-5* and *pif457* lines together with a wild-type (Col-0) control were germinated and grown under W for one week on plates at two different densities: low density (LD) and high density (HD). While WT plants were significantly longer in the HD setting, the length of *sav3-5* or *pif457* hypocotyls was similar under LD and HD (Figure 5), confirming that auxins are required for enhanced growth under high planting densities.

Open field trials with tomato were then carried out using the same pool of M82 and IL2-2 seeds in two different locations: Acerra (Campania, Italy) and Paiporta (Valencia, Spain). Plants were germinated in the greenhouse and transferred to the field after 3 weeks, at the end of April 2024. They were grown in rows separated 1 m, at two different densities: 6 plants per m^2^ (referred to as low density, LD) or 12 plants per m^2^ (high density, HD). The same row had two alternating groups of plants of each genotype (Supplemental Figure S3). About three months after planting, plants were analyzed and fruits were harvested and used for analysis (Figure 6). At the time of harvest, the plants from the Italian location were smaller and had produced less and smaller fruit compared to those in Spain (Figure 6), most likely due to their different growth conditions (soil, climate, etc.).

**Figure 6.**
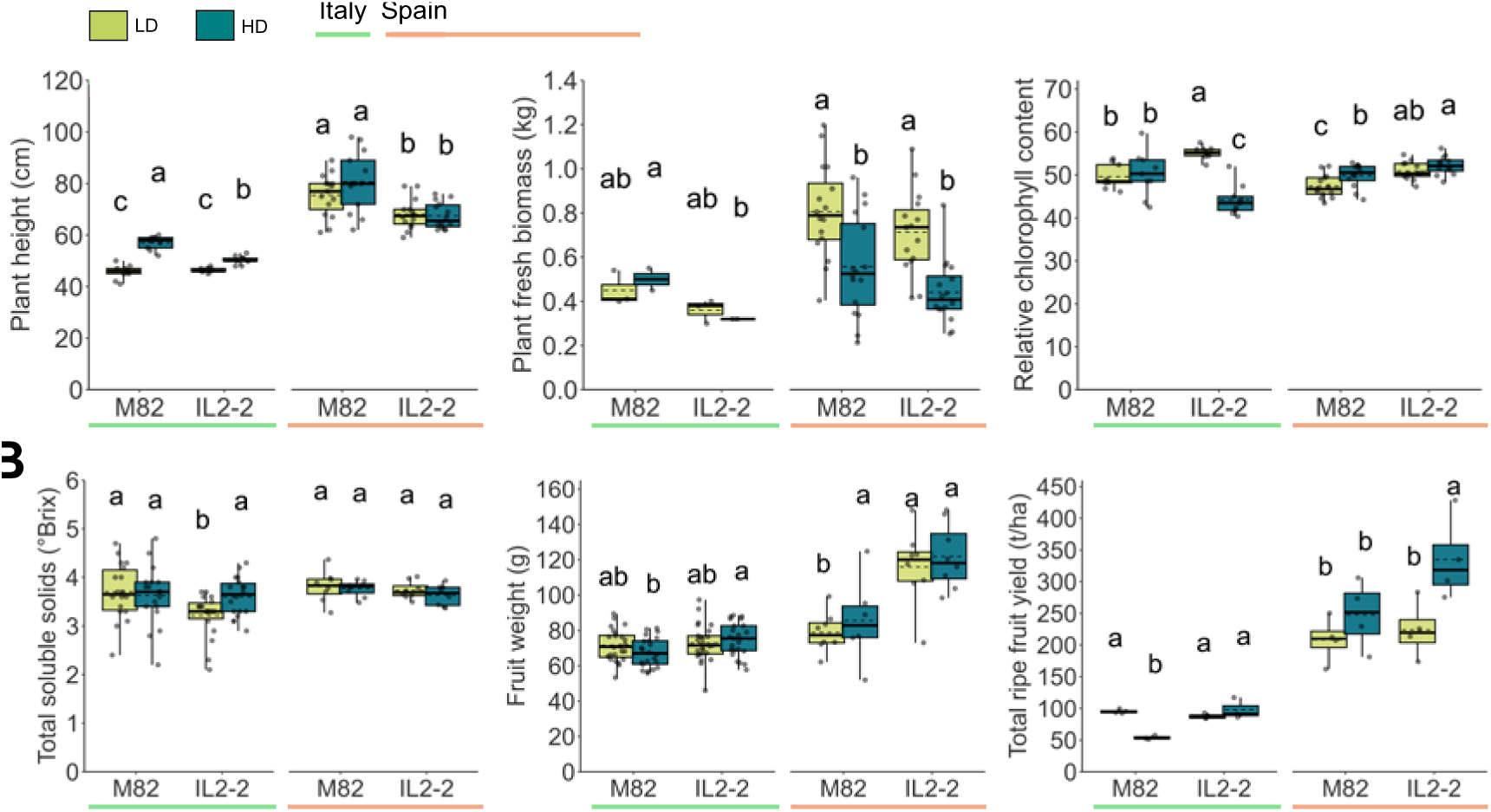
IL2-2 plants perform better than M82 under high planting densities. M82 and IL2-2 plants were grown in open fields located in Italy and Spain either at low density (LD) or high density (HD) conditions. (A) Plant height, biomass and leaf chlorophyll levels at the time of harvest. (B) Ripe fruit features and total yield. Boxplot legend and statistical analysis were as described in Figure 1 legend.

In both locations (Italy and Spain), M82 plants tended to be taller under HD conditions, as expected. Importantly, the difference in height under HD and LD conditions was attenuated in IL2-2 plants (Figure 6A), consistent with their auxin-related shade-tolerant phenotype. The effect of the HD conditions in the field also paralleled that of W+FR irradiation in the greenhouse in terms of plant biomass, that remained unchanged in all genotypes and conditions despite the described differences in plant height (Figure 6A). In the case of chlorophyll levels, however, the drop triggered by W+FR treatment (Figure 3B) was only observed when comparing IL2-2 plants grown under HD and LD in the Italian field (Figure 6A). Soluble content (°Brix) was similar in M82 and IL2-2 fruit from plants growing under LD or HD. Fruit weight was also unaffected by the plant genotype or the density of the culture in Italy but was higher in the IL2-2 line in Spain, particularly under HD conditions (Figure 6B). Most strikingly, fruit yield per hectare was similar in M82 and IL2-2 under LD conditions in both locations, but it was higher in the introgression line under HD (Figure 6B).

## Discussion

Tomato is a shade avoider plant that exhibits a pronounced multilevel response to proximity shade (low R/FR) as shown here and elsewhere (Cagnola et al., 2012; Chitwood et al., 2012; Bush et al., 2015; Chitwood et al., 2015;Fanwoua et al., 2019; Ji et al., 2019; Kim et al., 2019; Ji et al., 2020; Sun et al., 2020; Ji et al., 2021; Li et al., 2024; Shomali et al., 2024). In this work, we compared the most conspicuous phenotypic responses, i.e., enhanced growth and decreased accumulation of photosynthetic pigments, in different tomato accessions grown under the same experimental conditions (Figure 1). SAS in domesticated tomatoes is quite variable and attenuated compared to the strong shade avoidance displayed by wild tomato species (Chitwood et al., 2012; Bush et al., 2015). We confirmed that the domesticated accessions that we used for this work show similar leaf expansion and chlorophyll loss responses to low R/FR but a remarkable variation in stem elongation responses, being strongest in MoneyMaker and weakest in MicroTom (Figure 1). In particular, the epicotyl elongation response was the most reproducible and it was evident even in MicroTom, an accession that showed no significant hypocotyl elongation upon W+FR exposure in our experimental conditions (Figure 1). Based on these results, we carried out a screening for tomato lines with an attenuated epicotyl growth upon exposure to W+FR that led us to the identification of the *S. pennellii* introgression line IL2-2 (Figure 2 and Supplemental Figure S1). While this line showed a clear attenuation of shade-induced stem elongation in both seedlings and adult plants, the response to W+FR in terms of chlorophyll loss was almost undistinguishable from that of the parental line M82 (Figure 2 and Figure 4). Consistently, RNAseq data confirmed that IL2-2 plants show an impaired shade-triggered regulation of genes involved in growth (i.e., auxin-related) but not of those involved in photosynthetic pigment production (i.e., MEP, carotenoid, and chlorophyll biosynthetic pathways) (Figure 3 and Supplemental Figure S2). These results together confirm that the molecular pathways regulating elongation and photosynthetic pigment accumulation in response to proximity shade (low R/FR) in tomato are different and can be genetically separated, as previously shown in Arabidopsis (Toledo-Ortiz et al., 2010; Bou-Torrent et al., 2015; Morelli et al., 2021).

The perception and transduction of the low R/FR signal to regulate SAS responses is best understood in Arabidopsis (Martinez-García & Rodriguez-Concepcion, 2023). The light signal is sensed by phytochromes, with phyB playing the most relevant role in proximity shade. Under high R/FR, active phyB represses SAS responses by interacting with PIFs to repress their activity (Leivar & Monte, 2014). Members of the so-called PIF quartet (PIFQ: PIF1, PIF3, PIF4 and PIF5) and PIF7 directly contribute to the SAS (Martinez-García & Rodriguez-Concepcion, 2023). While both *pifq* and *pif7* mutants show a decreased elongation in response to W+FR, the *pif7* mutant shows a WT phenotype in terms of simulated shade-triggered photosynthetic pigment loss (Toledo-Ortiz et al., 2010; Bou-Torrent et al., 2015; Morelli et al., 2021). Because PIF1 and PIF3 are more associated to the repression of photosynthetic development whereas PIF4 and PIF5 are more related to the promotion of growth, it is possible that the molecular mechanism responsible for the shade tolerance phenotype of IL2-2 (affecting elongation growth but not photosynthetic pigment levels) utilizes signal transduction pathways involving PIF4, PIF5 or/and PIF7. These particular PIFs promote shade-triggered growth by preferentially regulating genes involved in the biosynthesis and transport of hormones, with a prominent role of the auxin IAA (Iglesias et al., 2018; Casal & Fankhauser, 2023; Martinez-Garcia & Rodriguez-Concepcion, 2023). IAA synthesized in shade-exposed cotyledons through the TAA/YUC pathway (Figure 5A) is transported to hypocotyls, where it stimulates cell elongation. Consistently, low R/FR regulates the expression of genes encoding PIN-FORMED (PIN) polar-auxin-transport efflux carriers. Upon arrival in hypocotyls, auxin binding to TRANSPORT INHIBITOR RESPONSE 1/AUXIN SIGNALING F-BOX (TIR1/AFBs) receptors leads to the degradation of the AUXIN/INDOLE-3-ACETIC ACID (AUX/IAA) proteins, which regulate the expression of genes for AUXIN RESPONSE FACTORS (ARFs). Other auxin-related genes rapidly induced by low R/FR include members of the *AUX/IAA* and the *SMALL AUXIN UP RNA* (*SAUR*) gene families, which are often upregulated in both cotyledons and hypocotyl tissues. Direct control of the expression of genes involved in both auxin-biosynthesis (e.g., *YUCs*), transport (e.g., *PINs*) and response (e.g., *IAAs*, *SAURs*) by Arabidopsis PIF4, PIF5, and PIF7 (Iglesias et al., 2018; Casal & Fankhauser, 2023) rapidly links the perception of low R/FR with activated plant growth.

In tomato, transcriptomic analyses have also found a central role for auxins in the response to low R/FR (Cagnola et al., 2012; Bush et al., 2015; Sun et al., 2020). We observed that genes related to auxin synthesis (*TAR1*/*TAA3*, *YUC5*, *YUC9*), transport (*PIN3*, *PIN4*), and signaling (*ARF10B*, *IAA1*, *IAA7*, *IAA17*, *IAA22*, *IAA29*, *SAUR*) show an altered expression in IL2-2 plants compared to the M82 parental in the absence of the low R/FR signal (Figure 3). W-grown IL2-2 plants also show increased expression of *BG1LA* and *BG1LE*, tomato homologs of the rice *BIG GRAIN 1* (*BG1*) gene whose overexpression increases auxin basipetal transport, auxin sensitivity, and grain size (Liu et al., 2015). Furthermore, two tomato homologs of the transcription factor ATHB2, a key early transducer of the low R/FR signal that reduces auxin responses (Iglesias et al., 2018; He et al., 2020), are also up-regulated in W-grown IL2-2 (Figure 3D). Interestingly, most of these genes show a reduced response to W+FR in the IL2-2 line, reaching transcript levels similar to those in shade-treated M82 plants (Figure 3). These observations suggest that IL2-2 plants have a strongly altered auxin homeostasis that hardly impairs growth or fruit development under high R/FR conditions but prevents a normal response to low R/FR (Figure 4, Figure 6, and Supplemental Figure S1). Further work should establish the molecular basis of this characteristic phenotype.

Despite most of the work on the participation of auxin on SAS responses has been done in lab conditions using W+FR to simulate proximity shade, our results using Arabidopsis *pif457* and *sav3* mutants grown at different densities confirmed that this hormone-dependent SAS pathway is also key to enhance growth under HD conditions (Figure 5). In agreement, the system-level rearrangement of auxin homeostasis deduced in IL2-2 plants (Figure 3) not only prevents normal elongation growth in response to FR supplementation in the greenhouse (Figure 2 and Figure 4) but also under HD conditions in the field (Figure 6). From the agronomic point of view, the IL2-2 line has several advantages. First, it is very similar to the M82 parental in terms of growth, biomass, photosynthesis, and fruit quality under high R/FR conditions, i.e., when grown under W (Figure 4) and LD (Figure 5). The only significant changes were detected in fruit hardness and seed number in the greenhouse, which were slightly decreased in IL2-2 (Figure 4D) and in leaf chlorophyll content and fruit weight in the field, which were increased (Figure 5). Second, it does not interfere with the beneficial effects that FR supplementation provides for fruit quality, i.e., increased content of soluble solids (°Brix), ascorbic acid (vitamin C) and carotenoids (pro-vitamin A) in ripe fruit (Figure 4D). And third, under HD conditions in the field it produced more fruit per area of cultivated land compared to the M82 parental (Figure 5B).

The work reported here demonstrates that identification of shade-tolerant tomato lines can be beneficial for agronomic practices involving high planting density. Our screening for shade-tolerant tomato lines used W+FR to simulate shade and epicotyl elongation as an output of the response (Figure 2 and Supplemental Figure S1). Implementing new screenings using HD instead of FR supplementation might identify new lines with improved performance in agricultural settings involving proximity shade. Another option would be to measure shade-triggered chlorophyll loss instead of epicotyl elongation as an output. Several methods exist for the high-throughput analysis of chlorophyll levels indoor or in open fields (Tayade et al., 2022), allowing to design large scale screenings. An added advantage of a chlorophyll-based screening is that the molecular mechanism responsible for the shade tolerance phenotype will likely be different from those responsible for the elongation-based phenotypes. Therefore, stronger lines could be creating by combining the two types of mechanisms (i.e., the two molecular pathways) causing the reduced response to low R/FR. We speculate that these lines could also perform better when intercropped with taller plants. Intercropping is a conservation agriculture practice that involves growing two or more crops in close proximity to one another. Intercropping promotes biodiversity and enhances crop resilience to extreme environmental changes, but it also increases crop yields in both low-input (traditional) and high-input (intensive) agrosystems (Tilman 2020; Li et al., 2023). However, development of intercropping is currently limited by the reduced toolbox of crop varieties amenable to this farming practice. Screenings for shade-tolerant lines or targeted manipulation of specific components of the low R/FR signal perception and transduction should provide a solution for this challenge.

## Acknowledgements

We thank the staff at the IBMCP Metabolomics Platform for technical support.

## Author contributions

EB-E, SF, JFM-G, MMR and MR-C conceived the project and designed the experiments; EB-E, SF, JP-R, AB, LV-C, JP-B and MA performed the experiments; EB-E, SF, JFM-G, MMR and MR-C analyzed and discussed the data; EB-E, SF, JFM-G, MMR and MR-C wrote the paper; all authors reviewed and approved the manuscript.

## Data availability

The data that support the findings of this study are available from the corresponding authors upon reasonable request.

## Supporting information

### SUPPLEMENTAL FIGURES

**Figure S1.**
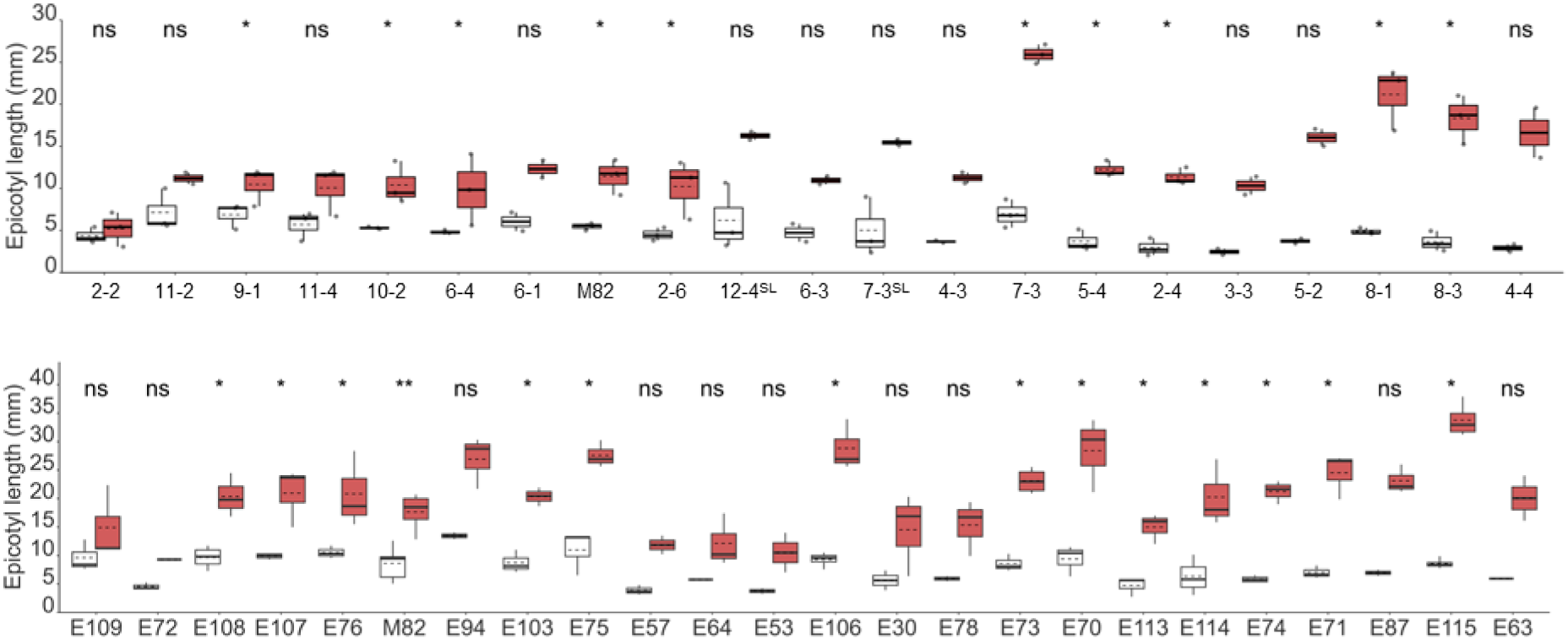
Differential responses of tomato accessions to simulated shade. Seeds from the indicated tomato lines were germinated and grown under white light (W) and then exposed to FR-supplemented W (W+FR) or left under W for 11 days. Values correspond to the length of the epicotyl after the W+FR (or control W) treatments. Dots represent individual data points. The lower boundary of the boxes indicates the 25th percentile, the black line within the boxes marks the median, the dotted line within the boxes marks the mean, and the upper boundary of the boxes indicates the 75th percentile of the data distribution. Whiskers above and below the boxes indicate the minimum and maximum values (excluding outliers). Statistically significant differences between W and W+FR values are represented with asterisks (t-test, ns: not significant; *: P < 0.05; **: P < 0.01).

**Figure S2.**
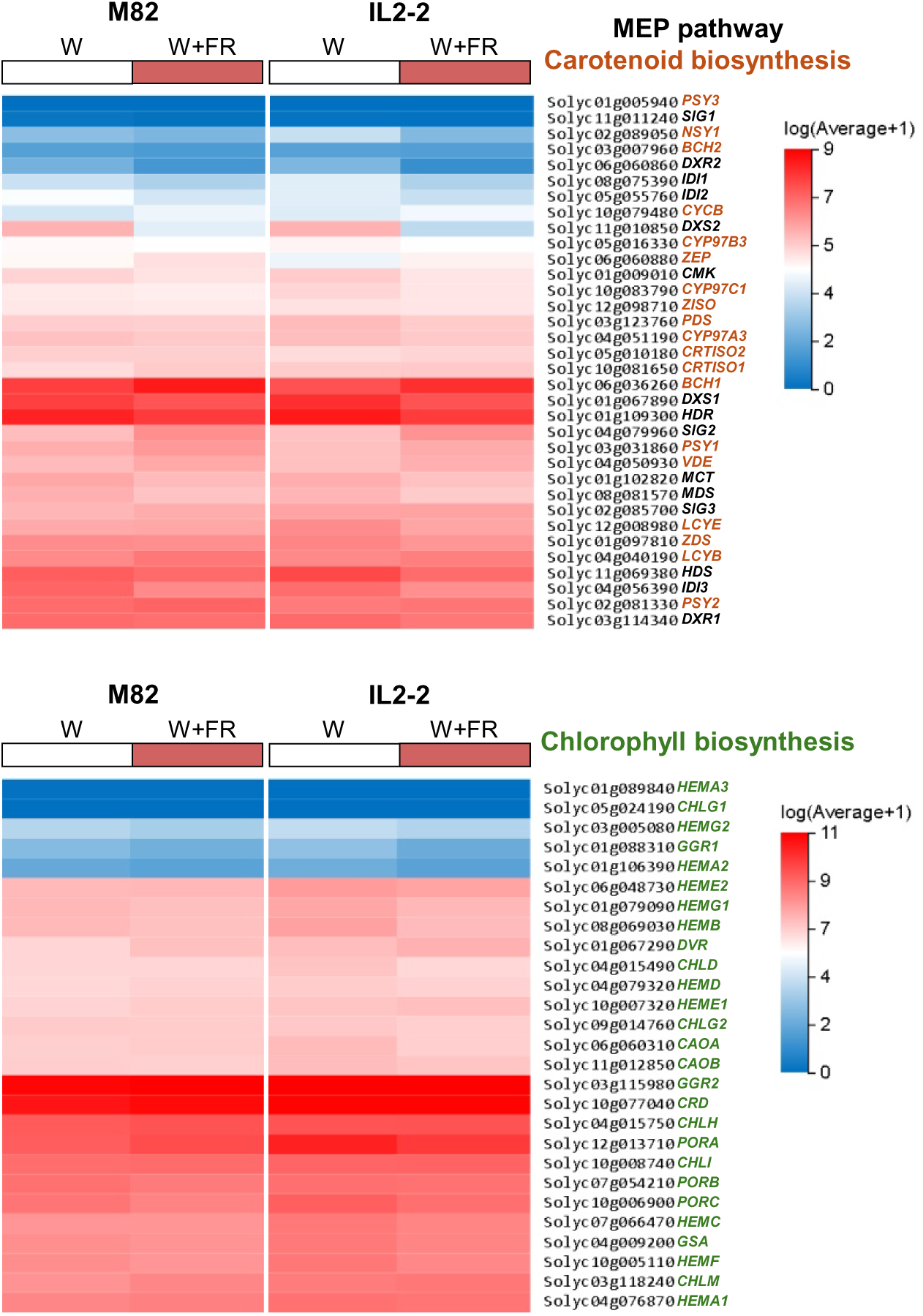
Heatmap comparison of the expression of genes involved in photosynthetic pigment biosynthesis. M82 and IL2-2 samples correspond to those described in Figure 3. Transcript abundance (mean log2 TPM+1) for the indicated genes

**Figure S3.**
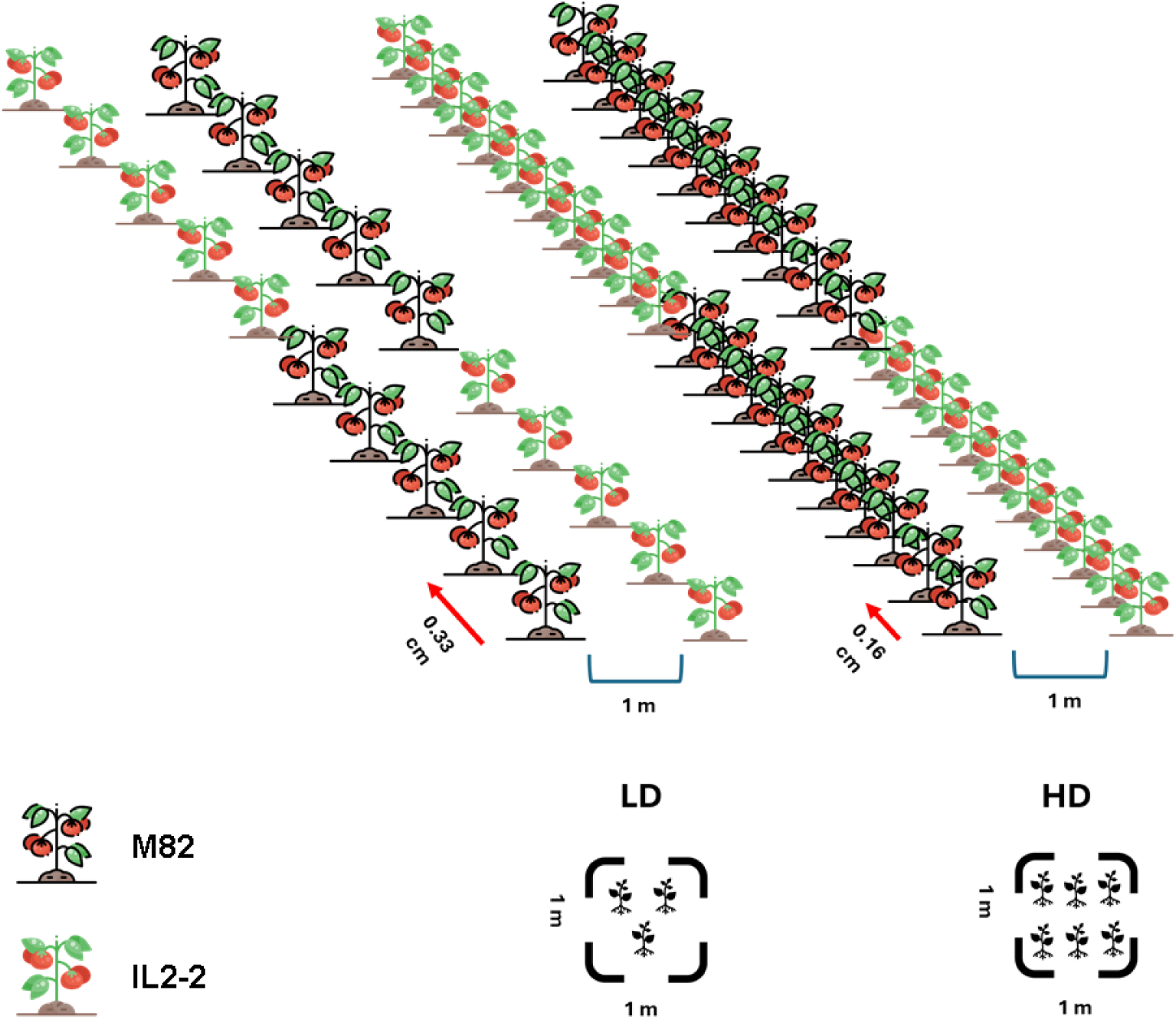
Schematic representation of the distribution of M82 and IL2-2 plants in the open field experiment. Number of plants per square meter under low density (LD) and high density (HD) conditions is also shown.

